# Physical activity interventions for major chronic disease: a matched-pair analysis of Cochrane and non-Cochrane systematic reviews

**DOI:** 10.1101/571901

**Authors:** Claudia Hacke, David Nunan

## Abstract

**Objective:** To assess the degree of concordance between Cochrane and non-Cochrane systematic reviews with meta-analyses of physical activity interventions.

**Study Design and Setting:** We conducted a matched-pair analysis with individual meta-analyses as the unit of analysis, comparing Cochrane reviews of randomised controlled trials of physical activity interventions with non-Cochrane reviews. Meta-analyses were matched based on the intervention, condition, outcome and publication year. Matched pairs were contrasted statistically in terms of differences between effect estimates, their precision, and number of citations using Wilcoxon two-sample test and agreement using Bland-Altman plots.

**Results:** Our search yielded 24 matched meta-analyses. Matched pairs were similar in terms of the number of included studies, sample sizes and publication year but only half (51.7%) of 545 individual clinical trials were included in both the Cochrane and non-Cochrane paired reviews. Effect estimates from non-Cochrane reviews were larger for 15 (62.5%) pairs, smaller for 8 (33.3%) and equal to Cochrane reviews for one (4.2%) pair. On average, effect estimates from non-Cochrane reviews were 0.12 log units (or 13%) higher compared with matched Cochrane reviews (z-score −2.312, P=0.012). We observed discrepancies with regard to the statistical (n=6) and clinical interpretation (n=4) of effect estimates, with non-Cochrane reviews reporting more often a statistically significant result (4/6) and effect sizes favouring intervention of greater than a two-fold (4/4) compared with Cochrane matches. Non-Cochrane reviews were also more frequently cited irrespective of whether the results agree or disagree in their statistical conclusion but this finding did not reach statistical significance at the traditional 0.05 threshold.

**Conclusion:** On average, meta-analyses from non-Cochrane reviews reported higher effect estimates and were more likely to show significant effects favouring the intervention compared with meta-analyses from Cochrane reviews. Though differences were small, they were sufficient to result in important discrepancies in statistical and clinical interpretations between a number of reviews.

**What is new?:** Key findings:

- Findings demonstrate non-Cochrane reviews on average report larger effect estimates and have discrepancies in statistical and clinical interpretation more likely to favour physical activity interventions. Potential sources underpinning discrepant review findings are explored in an accompanying sister paper.

What this adds to what was known?

- The first assessment of systematic differences between paired Cochrane and non-Cochrane meta-analyses examining the role of physical activity interventions for preventing and treating major chronic disease

What is the implication and what should change now?

- Authors should be aware of the need of protocol registration to minimise unnecessary duplication and be mindful of potential discrepant findings depending on the source of review evidence.

## 1. Introduction

Systematic reviews and meta-analyses are crucial instruments for informing evidence-based decisions in health care and underpinning guidelines and policy [1]. Their production has increased by more than 2500% since 1991, however, a high number of systematic reviews and meta-analyses have shortcomings and address the same topic with potentially redundant information [2-4].

Systematic reviews conducted via the Cochrane Collaboration are internationally recognized for their high methodological quality and rigor [5-7], Research evidence suggests that meta-analysis from systematic reviews published outside the Collaboration are of lower methodological and reporting quality, can provide discrepant results and conclusions, often demonstrating larger effect sizes and report more favourable conclusions [3, 4, 6, 8-11].

Two studies have addressed the issue by comparing pairs of meta-analysis of both types of reviews from pharmacological interventions using a matched-pair design [7, 8]. Jorgensen et al. summarized that Cochrane reviews compared with other industry supported meta-analyses of the same drugs in the same disease show almost similar estimated treatment effects, but the latter were found to be less transparent and reported more favourable conclusions without reservations about methodological limitations. More recently, Useem et al. [8] demonstrated that non-Cochrane reviews report larger effect sizes and with less precision than Cochrane reviews. The authors also noted that non-Cochrane reviews had a higher citation rate.

However, these two studies mainly considered pharmacological interventions in the field of cardiology and called for the need to replicate their findings in other fields and for other interventions. Importantly, as acknowledged in their limitations, neither study adequately addressed the underpinning reasons for the observed discrepancies between the reviews. We hypothesize that this problem likely concerns non-pharmacological treatments across many clinical areas, including lifestyle interventions such as physical activity and exercise-based interventions.

The aims of our study were therefore to explore whether systematic differences in findings exist between similar Cochrane and non-Cochrane reviews examining the role of physical activity interventions in prevention and management of major chronic disease. We also aimed to qualitatively explore the methodological reasons underpinning any observed discrepancies. In order to accommodate these two related but separate aims, we split our study in to two papers. In this paper we address quantitative treatment effect estimates reported in matched Cochrane and non-Cochrane reviews. In our sister paper we perform a qualitative assessment of both types of reviews in order to explore potential reasons for differences (Hacke & Nunan, 2019). The findings from both studies will be particularly relevant to policy makers and health care practitioners, ensuring any recommendations and shared decisions are informed by complete, accurate and high-quality evidence syntheses.

## 2. Material and Methods

### 2.1 Search, inclusion and matching strategies

An ongoing overview of systematic reviews assessing the effectiveness of physical activity interventions in preventing or treating major chronic diseases formed the basis of Cochrane review evidence in this study [12]. Accordingly, a literature search of the Cochrane Database of Systematic Review was undertaken to identify systematic reviews of RCTs assessing the effect of exercise-based interventions on clinical and patient relevant outcomes (search through December 2015). Inclusion was restricted to reviews relating to the 2008–2013 WHO non-communicable disease action plan including cardiovascular, respiratory, renal diseases, and cancer as well as metabolic, mental and musculoskeletal disorders.

A literature search for matching non-Cochrane reviews was then carried out using Google and Google Scholar followed by MEDLINE, TRIP Database, PEDro and Web of Science (see Appendix 1). Secondary searches involved hand-searching reference lists of relevant studies, expert guidelines, recommendations, reviews and meta-analysis.

Potentially eligible non-Cochrane meta-analyses were then matched to comparable Cochrane meta-analyses based on a three-step matching strategy:

#### 1. Match on the same *condition + intervention*

The first step was to search for matches that focussed on the same condition and intervention combination compared to the corresponding Cochrane review.

#### 2. Match on one of the primary *outcomes* of Cochrane review

Once matched on condition and intervention, potential non-Cochrane were checked for matching outcomes with the corresponding Cochrane review by two authors (CH and DN) and disagreements resolved by discussion or a third author (KM). To increase the likelihood of finding potential matches we checked all outcomes from non-Cochrane reviews with corresponding primary outcomes of the matched Cochrane reviews. Where no match on primary outcomes was found, we attempted to match on secondary outcomes.

#### 3. Match on the *year of publication*

After the above matching criteria were met, we selected the closest match to the Cochrane review according to the year of publication, restricted to within five years. When two or more potential matched non-Cochrane reviews were published in the same year, the one with the closest match on date of search was selected.

Thus, our approach ensures that each meta-analysis pair was as closely matched as possible in terms of the clinical question being answered and the period over which searches were performed. As a result we expected duplication and a high degree of overlap in the study results within matches, and where this was not found, reasons for discrepancies could be further explored.

### 2.2 Data extraction and quality assessments

One author (CH) extracted data and performed conversions and these were checked for accuracy by a second author (DN). Any discrepancies were resolved by discussion. The following data were extracted: author, title, journal, year of publication, disease condition, intervention (treatment), comparison (control), outcome, number and sample size of studies included, effect size, 95% confidence interval and heterogeneity statistics (e. g. I-squared). In our sister paper focusing on the qualitative evaluation of meta-analyses (Hacke & Nunan, 2019), we assessed the methodological quality of included reviews using AMSTAR (Assessing the Methodological Quality of Systematic Reviews) tool [13, 14]. In addition to AMSTAR appraisals, we extracted further bias-related reporting characteristics in more detail including the number of databases searched, search period, study design, publication status, language, list of included and excluded studies, errors in data extraction, QUOROM/PRISMA reference.

### 2.3 Statistical analyses

Review characteristics were summarized and displayed as frequency, median and interquartile range (IQR), or mean and standard deviation (SD). Differences in publication year, sample size and the number of studies included between Cochrane reviews and non-Cochrane reviews were calculated and tested for statistical significance using the Wilcoxon two-sample test.

Summary effect sizes for matched Cochrane and non-Cochrane paired meta-analyses were displayed in forest plots created in Microsoft Excel for visual comparison. In line with similar studies [8], we identified pairs with discrepant estimates, and categorized these based on the nature of the discrepancy as follows:

1. The statistical interpretation of the meta-analysis is discrepant due to differences in the width of 95% confidence interval (95% CI) around effect estimate, e.g., one review concludes a non-statistically significant effect and the other a significant effect;
2. The magnitude of effect sizes differed by at least 2-fold (but the effect is in the same direction);
3. The clinical interpretation of the meta-analysis differs due to differences in the direction of the effect size, e.g., one review favours intervention and the other favours control).

Furthermore, we quantified the degree to which summary effect and precision (95% CI) estimates differed between matched reviews using scatter plots on a logarithmic scale for visual illustration. Statistical differences in the average of reported effect sizes and precision estimates were analyzed using Wilcoxon two-sample test and agreement was assessed using Bland-Altman plots [15]. Statistical analyses were performed using IBM SPSS Statistics 23 (Chicago, IL).

If a matched-pair differed in unit of analysis for pooled estimates (e. g. mean difference versus standardised mean difference, relative risk versus absolute risk), we recalculated and converted the pooled effect estimate of the non-Cochrane meta-analysis. Where data for primary studies were not presented we also converted the non-Cochrane estimate to match with the estimate reported in the Cochrane review. As dichotomous and continuous outcomes needed to be combined in one pair, we re-expressed the odds ratio as a standardised mean difference using formula provided in the Cochrane Handbook [16]. All conversions were performed in RevMan (Version 5.3)[17] and we documented the number of cases in which conversions were performed.

Finally, we determined the rate of citation of each Cochrane and non-Cochrane review (until 19^th^ December 2017) using Google Scholar’s search engine. Citation rates were grouped by Cochrane and non-Cochrane reviews based on the nature of concordance and visually displayed by box plots.

## 3. Results

We identified 56 Cochrane reviews within the Cochrane Library and 108 potentially matched non-Cochrane reviews from the pre-specified database search (Fig. 1). Following full text screening and removal of reviews due to language restrictions our search and matching process yielded 24 matched pairs across six disease areas as follows:

**Fig. 1.**
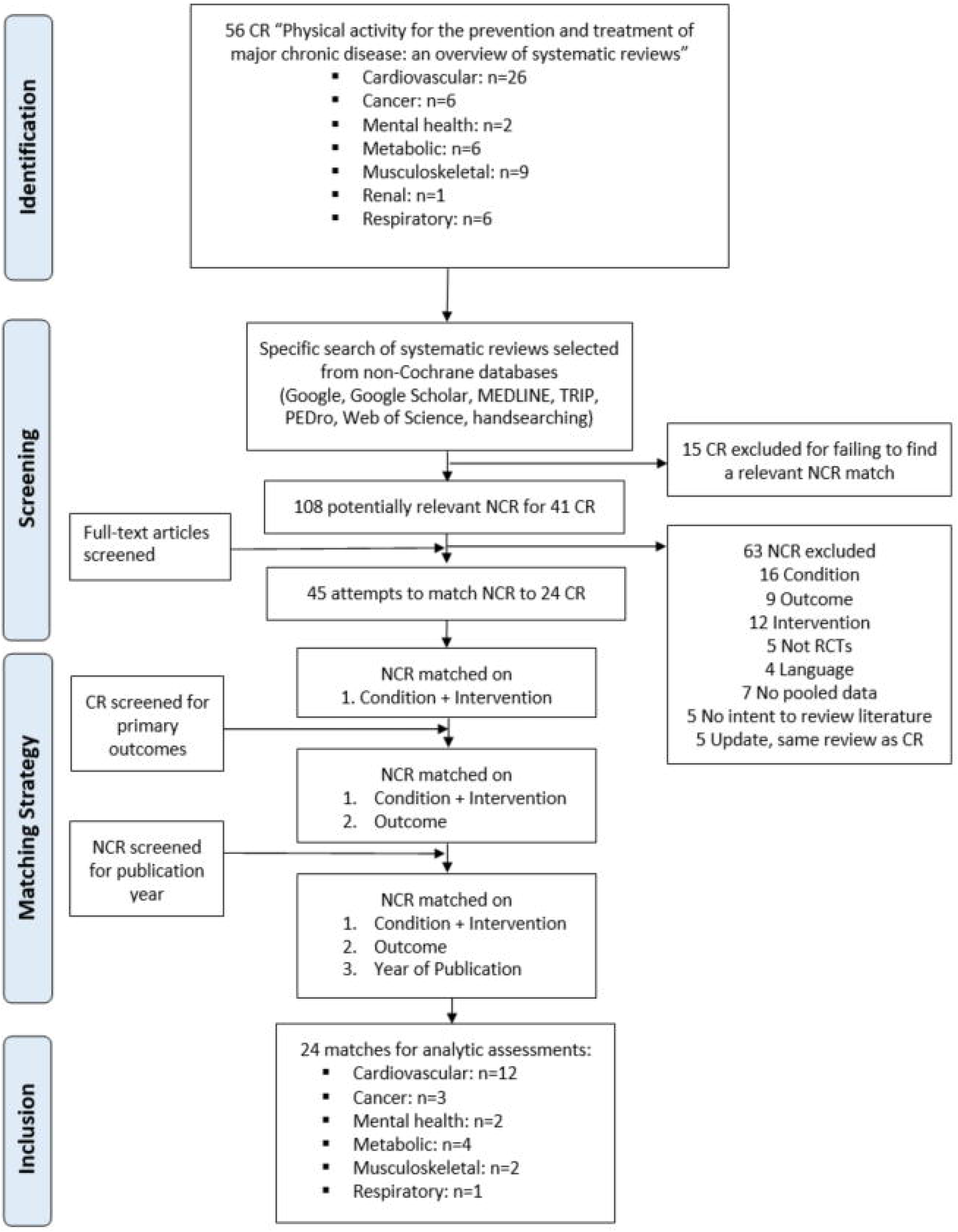
Flowchart of study selection and matching strategy. CR, Cochrane Review, NCR, Non-Cochrane Review

- 12 Cardiovascular diseases: 8 stroke, 1 heart failure, 2 intermittent claudication, 1 cardiac rehabilitation
- 3 Cancer: 1 breast cancer, 2 breast and prostate cancer
- 2 Mental health: 1 depression, 1 dementia
- 4 Metabolic: 3 diabetes, 1 obesity
- 2 Musculoskeletal: 2 osteoarthritis
- 1 Respiratory: 1 COPD

Of the 24 pairs, 5 pairs included the same non-Cochrane review as the matching study.

The majority of non-Cochrane reviews were excluded because they did not match on the same condition (28%) or intervention (23%). Another 7 non-Cochrane reviews reported that meta-analysis was inappropriate, whilst Cochrane review matches with pooled data were available. For two Cochrane reviews, a match could only be identified based on secondary outcomes.

### 3.1 Characteristics of included reviews

Characteristics of the included matched-pair reviews are shown in the supplementary table 1 (see Appendix 2). Overall, the 24 Cochrane reviews included a total of 259 studies and a total of 20,556 participants compared with 286 studies and 24,189 participants in the 24 non-Cochrane reviews. Of the 545 RCTs, 282 studies (51.7% in total) were found and included in both Cochrane and non-Cochrane reviews; 118 RCTs (22.2%) were found and included exclusively in Cochrane reviews and 145 studies (27.3 %) in non-Cochrane reviews. Taken together, half of the 545 RCTs were only included either in the Cochrane or in the non-Cochrane meta-analyses.

Cochrane reviews had a median of 7.5 (IQR 3.3-14.5) studies with a median sample size of 364 participants (IQR 215-1165). Non-Cochrane reviews had a similar total number of studies (median 7.5, IQR 6.0-13.0; p=0.340) and sample size (median 372 participants, IQR 243-1052; p=0.130). Across matched-pairs there was no difference in the how frequently one type of review was published prior to the other (p=0.434); Cochrane meta-analyses were published prior to its non-Cochrane match in 10 of 24 pairs, while publication of 8 non-Cochrane reviews were before the matched Cochrane review. Six matched pairs were published in the same year. The median publication year was 2013 (IQR 2010-2014) and 2014 (IQR 2010-2014) for Cochrane and non-Cochrane reviews, respectively.

### 3.2 Effect estimate comparisons

Figures 2 to 5 illustrate the comparison of summary effect sizes for matched Cochrane and non-Cochrane paired meta-analyses (n=24). The effect estimates with its corresponding 95% CI were recalculated and converted into the pooled effect estimate of its match in five Cochrane reviews (MD converted to SMD, OR converted to SMD) and three non-Cochrane reviews (SMD to MD, RD to RR). In this regard, we identified errors in the pooled effect estimate of one non-Cochrane review (pair 19). Overall, non-Cochrane reviews had larger effect sizes for 15/24 (62.5%) pairs, smaller for 8/24 (33.3%) and equal for 1/24 (4.2%).

**Fig. 2.**
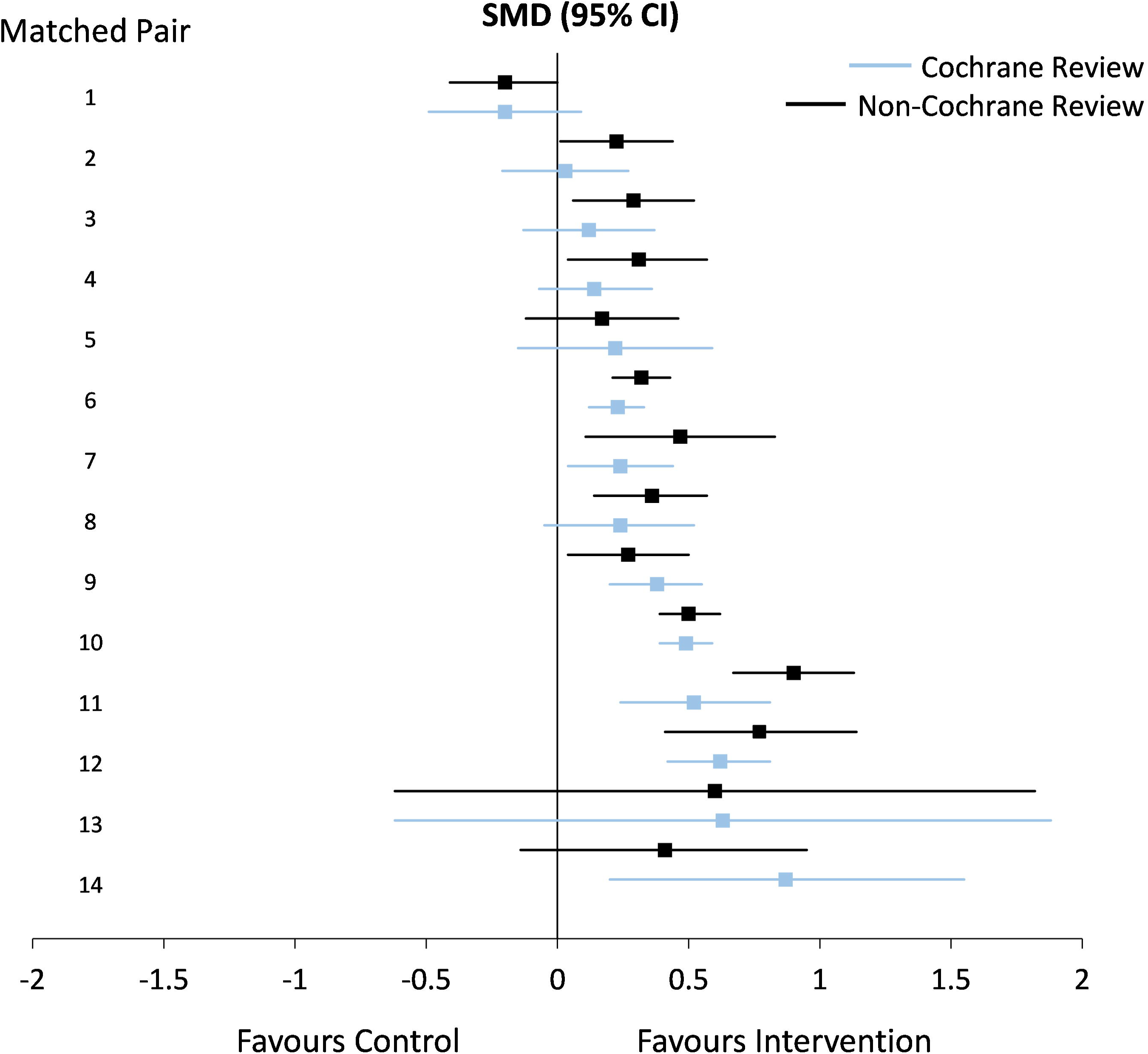
Summary effect estimates for matched Cochrane and non-Cochrane paired reviews. The figure illustrates a forest plot of effect sizes expressed as standardised mean differences with 95% confidence intervals for each pair of Cochrane (blue) and non-Cochrane (black) reviews. The matched pairs has been sorted based on effect size from the Cochrane review in ascending order

**Fig. 3.**
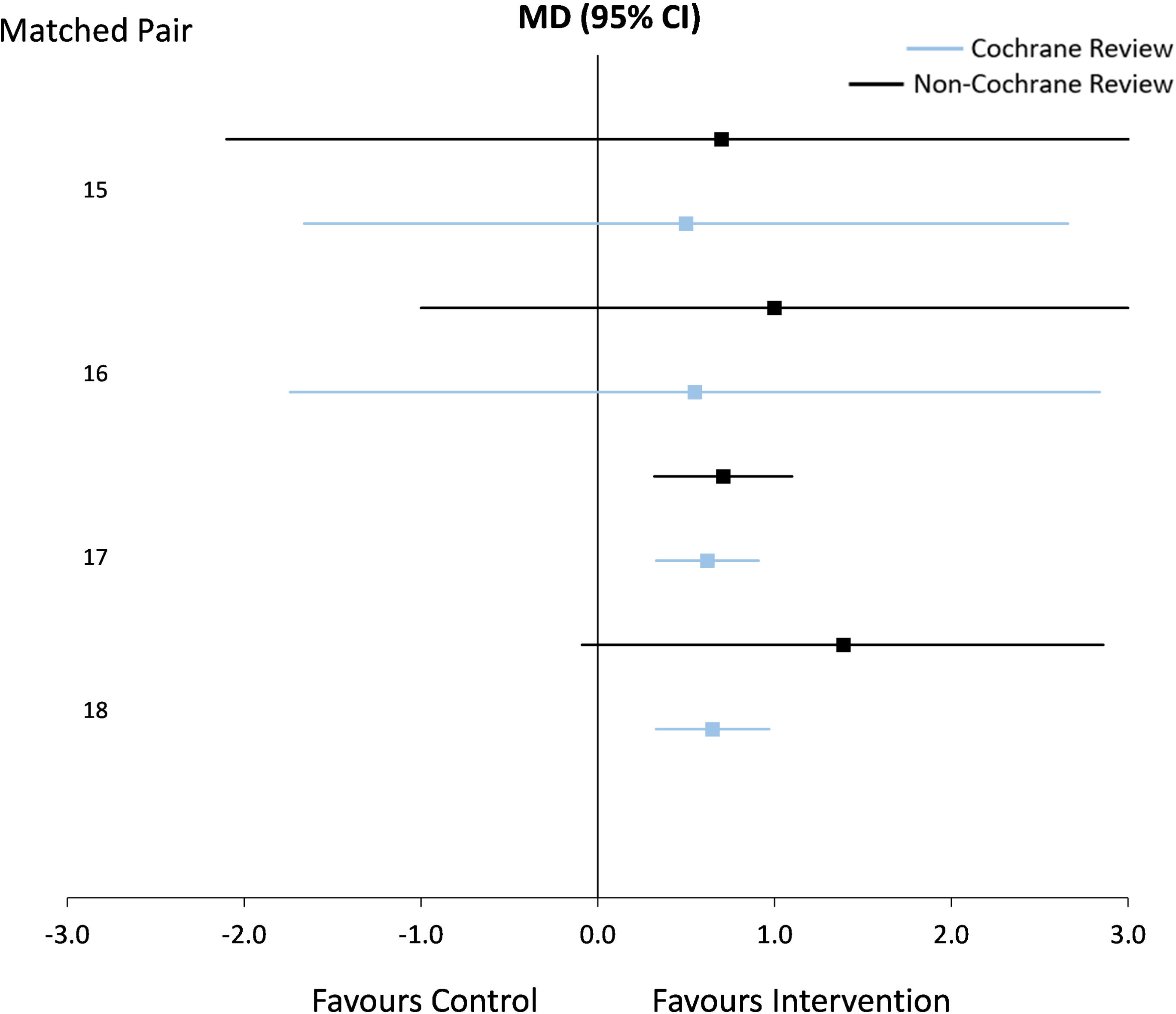
Summary effect estimates for matched Cochrane and non-Cochrane paired reviews. The figure illustrates a forest plot of effect sizes expressed as mean differences with 95% confidence intervals for each pair of Cochrane (blue) and non-Cochrane (black) reviews. The matched pairs has been sorted based on effect size from the Cochrane review in ascending order

The scatter plots of the summary effect estimates and their precision (95% CI) are displayed on a logarithmic scale in Figure 6. The Wilcoxon signed rank test showed that there was a significant difference between effect estimates reported in Cochrane and non-Cochrane reviews (z=-2.312, P=0.012). On average, effect estimates from non-Cochrane reviews were 0.12 log units higher, representing an average inflation of 13% (Figure 7). Expressed in original units, non-Cochrane reviews give larger estimates for SMD of 0.06 units and RR/HR of 0.26 units compared with their matched Cochrane review. There was no statistical difference in precision estimates of pooled effect estimates between reviews (z=-0.400, P=0.689).

**Fig. 4.**
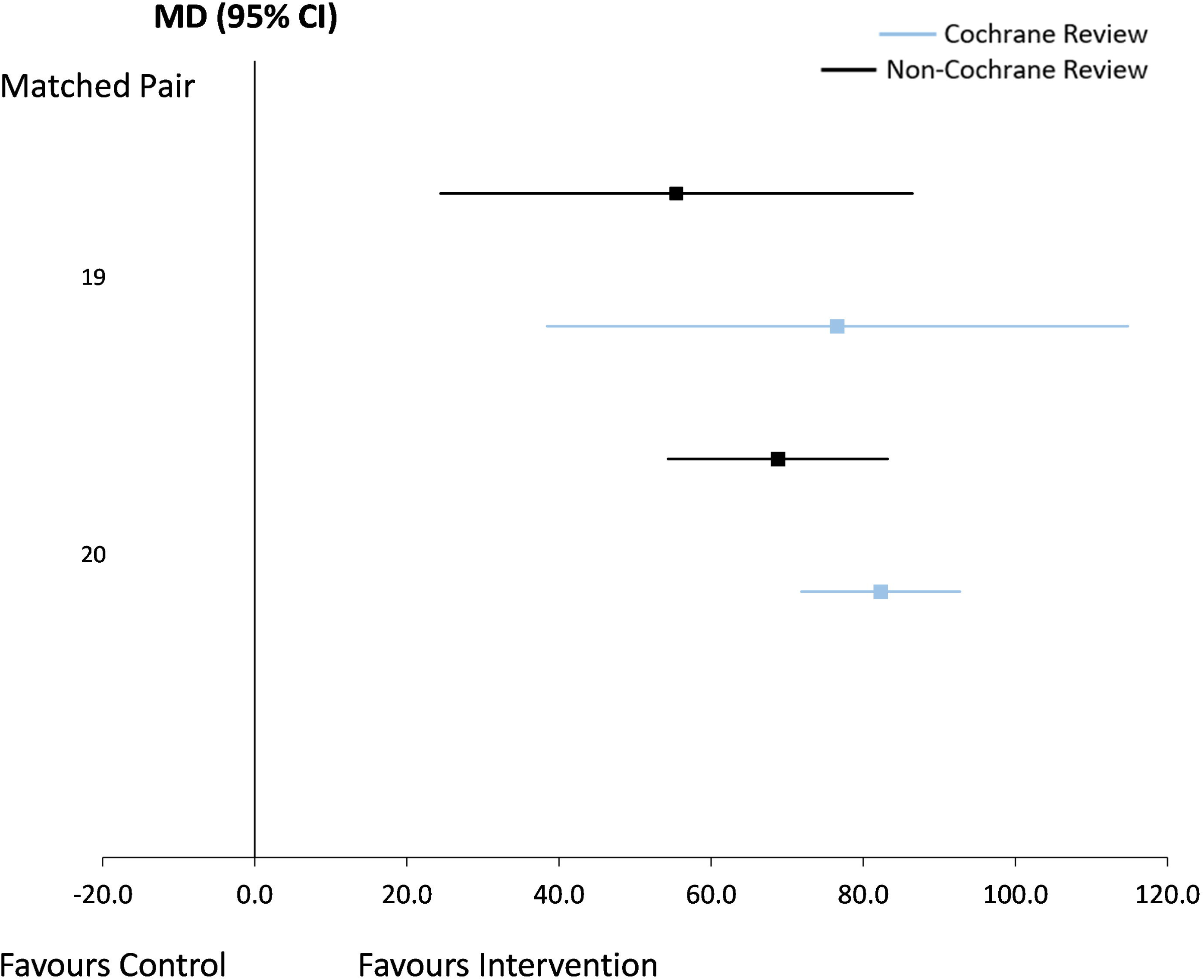
Summary effect estimates for matched Cochrane and non-Cochrane paired reviews. The figure illustrates a forest plot of effect sizes expressed as mean differences with 95% confidence intervals for each pair of Cochrane (blue) and non-Cochrane (black) reviews. The matched pairs has been sorted based on effect size from the Cochrane review in ascending order

**Fig. 5.**
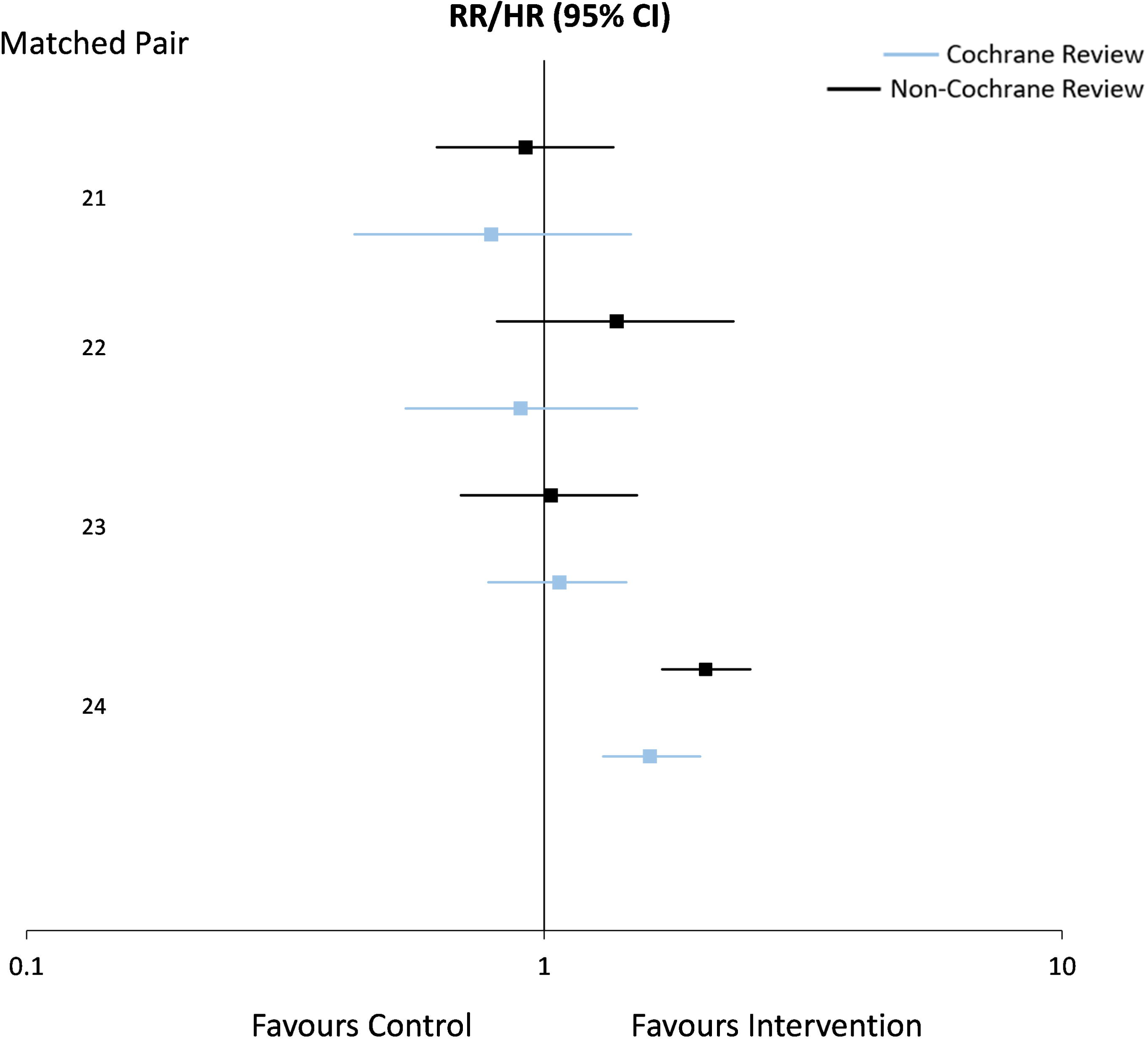
Summary effect estimates for matched Cochrane and non-Cochrane paired reviews. The figure illustrates a forest plot of effect sizes expressed as risk ratio/hazard ratio with 95% confidence intervals for each pair of Cochrane (blue) and non-Cochrane (black) reviews. The matched pairs has been sorted based on effect size from the Cochrane review in ascending order

**Fig. 6.**
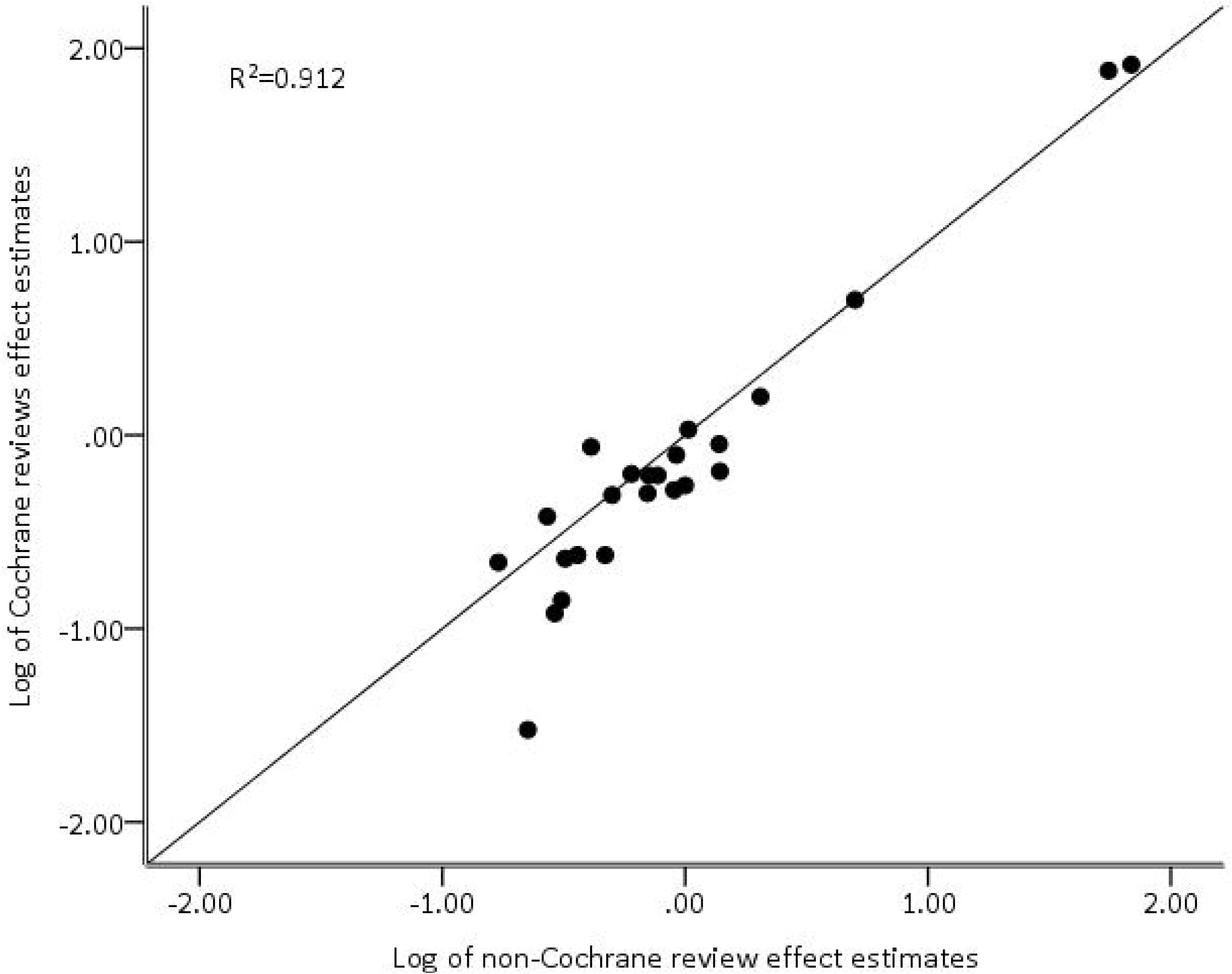
Scatter plot with line of equality of effect estimates from Cochrane and non-Cochrane matched metaanalyses in natural log units

**Fig. 7.**
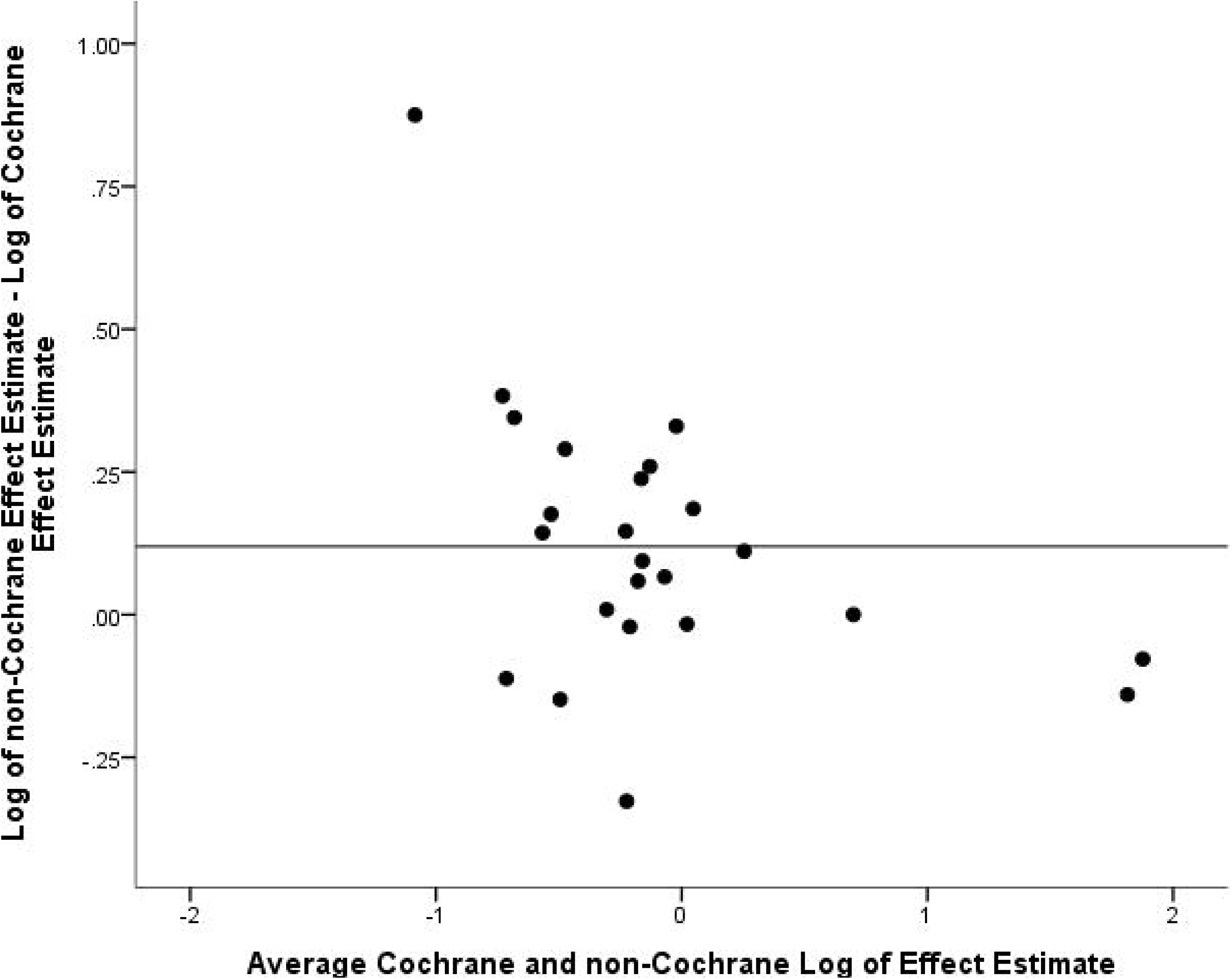
Bland-Altman plot showing the difference against the average of log-transformed non-Cochrane and Cochrane effect estimates

In six of the 24 matched pairs (25.0%) there were discrepancies in statistical interpretation due to differences in the width of 95% CI around the point estimate, with a statistically significant effect in one review but not in its match (Table 1). In four of these pairs, the non-Cochrane review reported a significant effect where the Cochrane review did not. Discrepancies in the direction of the effect estimate were observed in one of the 24 pairs. In this case the non-Cochrane review showed a nonsignificant protective effect favouring the intervention in contrast to the non-Cochrane review which showed a harmful effect. A 2-fold difference in the magnitude of effect estimate was observed in four pairs (18.2%), with the non-Cochrane review reporting the larger estimate in all cases.

**Table 1.**
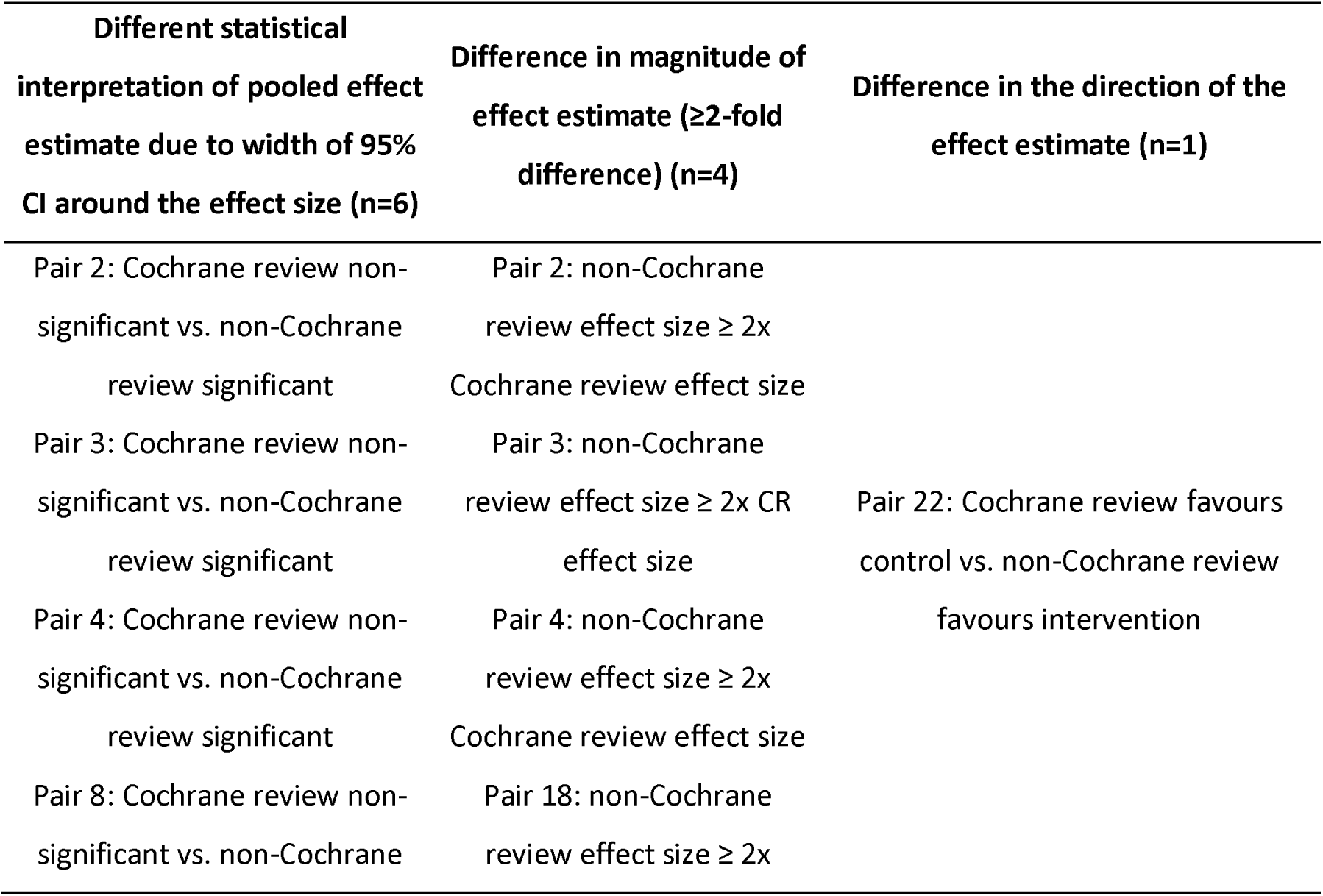
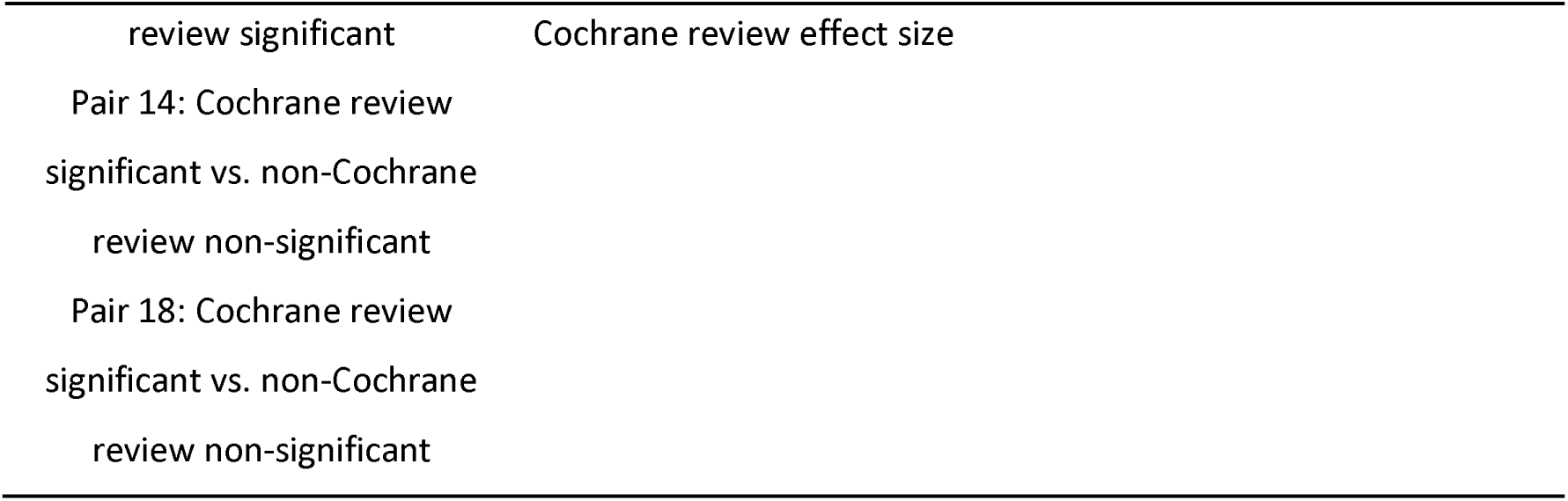
Summary of discrepant results between matched Cochrane and non-Cochrane reviews (24 pairs)

### 3.3 Study overlap across matched-pairs

Of the six matched-pairs coming from reviews published in the same year, 43 of 146 RCTs (58.9%) were included in both review types; 13/146 (8.9%) were included only in the Cochrane review and 47/146 (32.2%) only in non-Cochrane matched reviews (Table 2). Stated otherwise, despite reviews being published around the same time, there was a notable lack of overlap of included studies. Similar discrepancies in study overlap were found when the Cochrane or non-Cochrane review was published first, with only around half included in both reviews. Reasons for these discrepancies are explored in our sister paper (Hacke & Nunan, 2019).

**Table 2.**
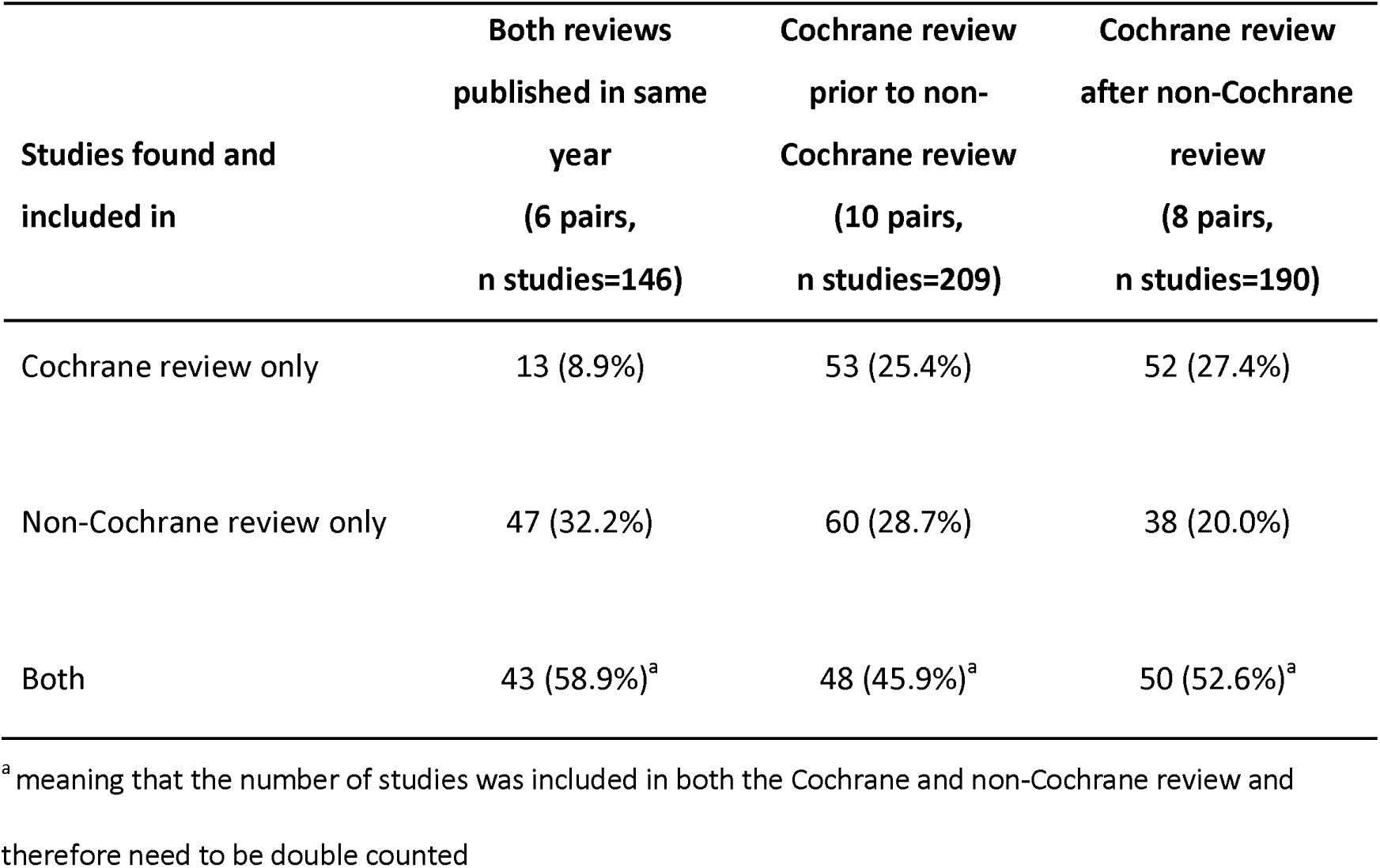
Overlap of studies by publication sequence

### 3.4 Citation rate

The number of times each type of review in a matched-pair was cited based on the nature of discrepancy is presented in Figure 8. Across all categories, median citation rates were higher in non-Cochrane reviews but statistically non-significant different from Cochrane review matches. The largest deviation was found when the discrepancy was due to shifts in 95% CI around the effect estimates resulting in conflicting statistical interpretations. Accordingly, non-Cochrane reviews were cited 264 times in the literature, while Cochrane reviews were cited 165 times, even though Cochrane reviews were published on average almost 2 years before the non-Cochrane matches. In other words, statistical significant and positive results were more likely to be cited by other articles given that in 4/6 of those pairs with discrepant statistical conclusions the non-Cochrane review reported the significant effect.

**Fig. 8.**
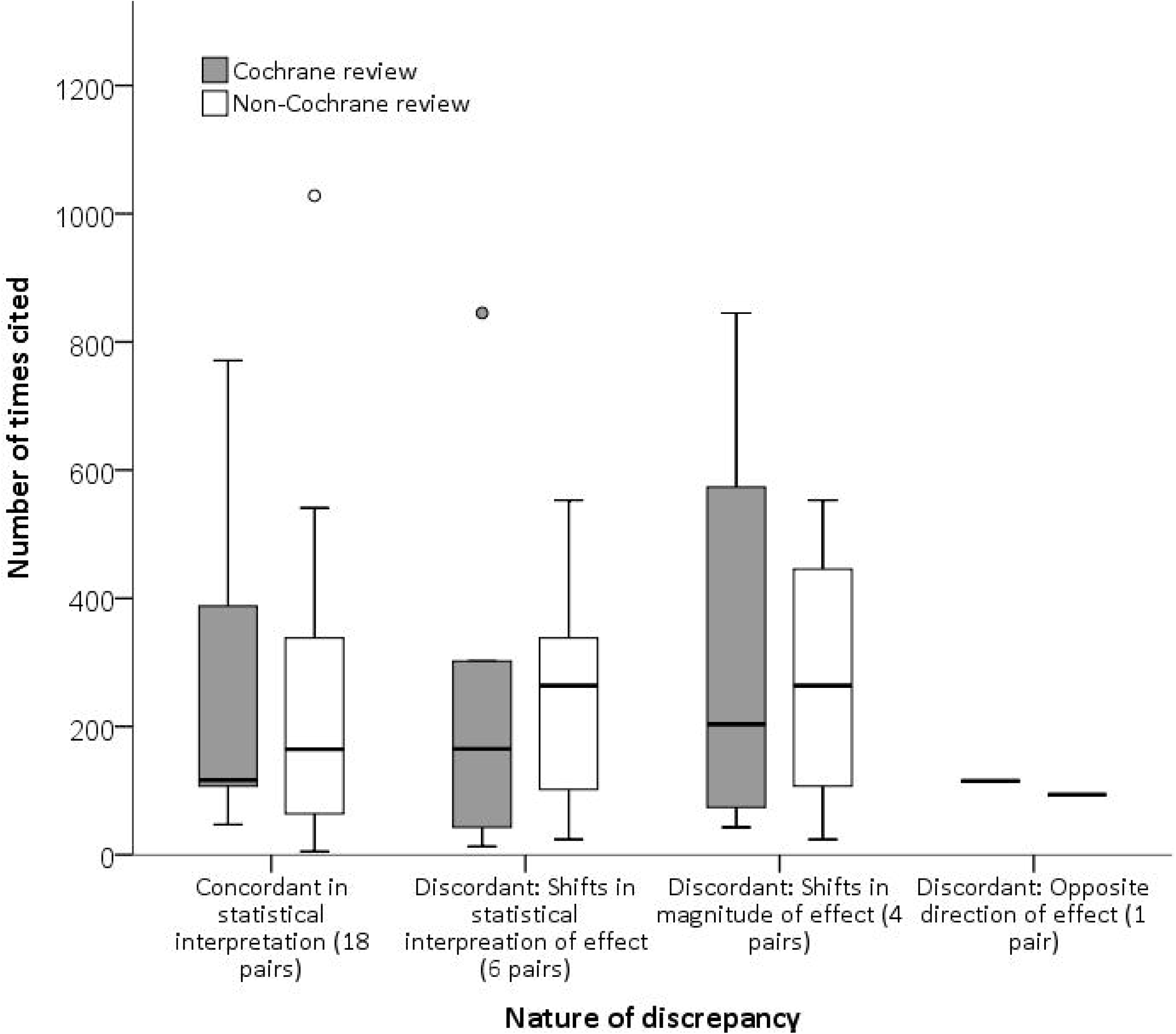
Citation rates of Cochrane and non-Cochrane matched pairs based on the nature of discrepancy in reported effect estimates

## 4. Discussion

### 4.1 Principle findings

This study is the first to evaluate differences between pooled effect estimates reported in Cochrane and non-Cochrane reviews asking similar questions of a non-pharmacological intervention. Moreover, our sister study is the first to provide a comprehensive assessment of methodological quality and offer potential explanations for differences in reported treatment effect estimates between the different types of reviews (Hacke & Nunan, 2019).

The main findings of this study were that non-Cochrane reviews reported larger, and occasionally clinically meaningful, effect estimates than matched Cochrane reviews. The most common discrepancy related to the statistical interpretation of the meta-analyses, where non-Cochrane reviews were more likely to report a statistically significant effect favouring the intervention. Non-Cochrane reviews were also more likely to be cited in the medical literature, even when the results of the contrasted reviews agreed in their statistical conclusion. Potential sources for the identified discrepancies are discussed in detail in our sister paper (Hacke & Nunan, 2019) and included differences in methodological quality, data abstraction procedures and disagreements in the interpretation of pre-defined eligibility criteria.

### 4.2 Comparison with other studies

We are aware of only two studies that have directly compared Cochrane and non-Cochrane reviews answering the same or similar question and using a matched pair design – focusing on pharmacological therapy [7, 8]. Our findings reinforce previous data that in general inconsistencies in treatment effect estimates between Cochrane and non-Cochrane reviews exist and cannot simply be explained by publication sequence [8]. The small, clinically insignificant difference of 0.12 log units observed for standardised effect estimates was surprising, particularly given the poor overlap of studies between Cochrane and non-Cochrane reviews (51.7%). This suggests that for standardised measures (average difference of *d=*0.06), results from Cochrane and non-Cochrane are likely to be congruent. However, this has to be balanced against almost one-quarter of matched pooled effect estimates with discrepancies that resulted in opposite statistical or clinical interpretations. A key question is what mechanisms are responsible for such discrepancies among a topic-matched pair. These are explored in our sister paper (Hacke & Nunan, 2019).

It is noteworthy that non-Cochrane reviews were cited more often in the medical literature across all of the given categories of discordancy. In light of the robust methodology and stringent guidelines for Cochrane reviews, it is somewhat surprising that this was also the case even when the results of the contrasted reviews agreed in their statistical conclusion. This may partly be explained by the fact that in the majority of these pairs (10/18 pairs) the non-Cochrane meta-analysis reported larger effect sizes than its matched Cochrane review. Accordingly, our findings supports further evidence of citation bias favouring significant study results in medical research [18-20].

Our analyses contribute to recent debates concerning research waste and redundancy due to the high prevalence of overlapping meta-analyses published on the same topic, reflecting wasted efforts and inefficiency [4, 21]. Collectively, these findings suggest several courses of action for improved coordination between reviewers and standardised methods of reporting, better registration of protocols and performing more inclusive publications and designs of systematic reviews, to minimise unnecessary replication across all fields of health care interventions.

### 4.3 Strengths and limitations of this study

A strength of our study is the effort made to ensure included reviews were matched as closely as possible on key elements including condition, intervention, and outcome. In addition, our approach to match on the year of publication helped minimise the probability of discrepant results due to publication sequence. In contrast to previous studies, we included both dichotomous and continuous outcomes and utilised appropriate statistical analysis methods to assess both differences and agreement between matched-pair pooled effect estimates.

The major limitation of this study is one that is common to all studies of this type, that being the potential for error due to reliance on secondary data as extracted and reported in published metaanalyses. Indeed, we report a large number of discrepancies concerning the extracted data of primary studies in our sister paper (Hacke & Nunan, 2019) and other errors cannot be ruled out. Accordingly, an issue that is not addressed in our study, nor previous studies, is which of the two sets of reviews more accurately extract and report data from the included primary studies. A greater focus on the correctness of extracted data in future comparisons could help further explain observed discrepancies between the two review types. Another limitation is the reliance on topics and conditions by Cochrane reviews as the evidence base for matching. We have therefore likely missed inclusion of other conditions for which there is a non-Cochrane review that was not covered in the Cochrane Database of Systematic Reviews.

## 5. Conclusion

Meta-analyses from Cochrane and non-Cochrane systematic reviews assessing the effect of physical activity interventions in chronic disease show discrepancies in pooled effect estimates and their statistical and clinical interpretation. Non-Cochrane reviews reported larger effect estimates and were more likely to favour the intervention when findings were discrepant between reviews. Awareness of the need for protocol registration could help minimise unnecessary duplication and avoidable research waste. Users applying systematic review evidence to inform evidence-based decisions around physical activity therapy need to be mindful of potential discrepant findings depending on the source of systematic review even if methodology rigorous.

## Funding

This research did not receive any specific grant from funding agencies in the public, commercial, or not-for-profit sectors.

## Declarations of interest

None.

## Data Availability

All data generated or analysed during this study are included in this published article [and its supplementary information files].

## Authors’ contributions

**C.H.; D.N.:** Conceptualization, Methodology, Validation. **C.H.:** Formal Analysis, Data curation, Investigation, Writing-Original draft preparation, Visualization. **D.H.:** Supervision. **C.H.; D.N.:** Writing-Reviewing and Editing.

## Acknowledgements

The authors would like to thank Nia W. Roberts for her role in the original search of Cochrane Systematic Reviews.

## Supplementary Material

**Appendix 1.** Search strategy

**Appendix 2.** Review characteristics

