## Appendix 1 for "Physical activity interventions for major chronic disease: a matched-pair analysis of Cochrane and non-Cochrane systematic reviews"

**Appendix 1. Search Strategy**

1. “(same title as Cochrane review)”, “systematic review”, “meta-analysis”

1. filter: published within same year of Cochrane

2. filter: published within 5 years above and below Cochrane review

2. “(name of intervention)”, “(condition of CR)”, “systematic review”, “meta-analysis”

1. filter: published within same year of Cochrane

2. filter: published within 5 years above and below Cochrane review

3. “exercise” “activity” “physical fitness” “rehabilitation” “sports therapy” “recreation”, “(condition of CR)”, “systematic review”, “meta-analysis”

1. filter: published within same year of Cochrane

2. filter: published within 5 years above and below Cochrane review
